# A meta-analysis of immune cell fractions at high resolution reveals novel associations with common phenotypes and health outcomes

**DOI:** 10.1101/2023.03.20.533349

**Authors:** Qi Luo, Varun B. Dwaraka, Qingwen Chen, Huige Tong, Tianyu Zhu, Kirsten Seale, Joseph M Raffaele, Shijie C. Zheng, Tavis L. Mendez, Yulu Chen, Sofina Begum, Kevin Mendez, Sarah Voisin, Nir Eynon, Jessica A. Lasky-Su, Ryan Smith, Andrew E. Teschendorff

## Abstract

**Background:** Changes in cell-type composition of complex tissues are associated with a wide range of diseases, environmental risk factors and may be causally implicated in disease development and progression. However, these shifts in cell-type fractions are often of a low magnitude, or involve similar cell-subtypes, making their reliable identification challenging. DNA methylation profiling in a tissue like blood is a promising approach to discover shifts in cell-type abundance, yet studies have only been performed at a relatively low cellular resolution and in isolation, limiting their power to detect these shifts in tissue composition.

**Methods:** Here we derive a DNA methylation reference matrix for 12 immune cell-types in human blood and extensively validate it with flow-cytometric count data and in whole-genome bisulfite sequencing data of sorted cells. Using this reference matrix and Stouffer’s method, we perform a meta-analysis encompassing 25,629 blood samples from 22 different cohorts, to comprehensively map associations between the 12 immune-cell fractions and common phenotypes, including health outcomes.

**Results:** Our meta-analysis reveals many associations with age, sex, smoking and obesity, many of which we validate with single-cell RNA-sequencing. We discover that T-regulatory and naïve T-cell subsets are higher in women compared to men, whilst the reverse is true for monocyte, natural killer, basophil and eosinophil fractions. In a large subset encompassing 5000 individuals we find associations with stress, exercise, sleep and health outcomes, revealing that naïve T-cell and B-cell fractions are associated with a reduced risk of all-cause mortality independently of age, sex, race, smoking, obesity and alcohol consumption. We find that decreased natural killer cell counts are associated with smoking, obesity and stress levels, whilst an increased count correlates with exercise, sleep and a reduced risk of all-cause mortality.

**Conclusions:** This work derives and extensively validates a high resolution DNAm reference matrix for blood, and uses it to generate a comprehensive map of associations between immune cell fractions and common phenotypes, including health outcomes.

**Availability:** The 12 immune cell-type DNAm reference matrices for Illumina 850k and 450k beadarrays alongside tools for cell-type fraction estimation are freely available from our EpiDISH Bioconductor R-package http://www.bioconductor.org/packages/devel/bioc/html/EpiDISH.html

## Background

Human tissues contain many different cell-types in proportions that vary between healthy individuals as well as in association with disease and exposure to disease risk factors [1, 2]. These shifts in cell-type proportions may not only constitute important biomarkers of environmental exposures, disease risk or early diagnosis, but may be causally implicated, as exemplified by immune-cell variations that impact cancer progression [3] and immuno-senescence [4]. Although detecting shifts in cell-type composition in easily accessible tissues like blood has been possible with moderately sized studies in the context of auto-immune diseases, cancer or aging [5-8], detecting more subtle changes in cell-type proportions that may arise in relation to disease risk factors like gender, obesity or smoking, has been more challenging. This is not only because underlying shifts in cell-type proportions may be of low magnitude, typically involving only a few percentage points, but also because there is already substantial variation in these proportions between healthy or unexposed individuals. Thus, measuring cell-counts in large cohorts of samples is necessary in order to confidently identify disease-or-exposure associated shifts in cell-type composition. However, experimental cell-counting methods are cumbersome and not easily scalable to thousands of samples.

DNA methylation (DNAm) has been abundantly profiled in easily accessible and heterogeneous tissues like whole blood [9-15], saliva [16, 17] and buccal swabs [18]. The underlying cell-type heterogeneity (CTH) of these tissues thus offers the opportunity to detect phenotype-associated shifts in cell-type composition. Indeed, because DNAm is highly cell-type specific and can be measured with high accuracy [19], application of cell-type deconvolution algorithms [2, 20] to average DNAm profiles generated by Epigenome-Wide-Association-Studies (EWAS) has proved to be an excellent means to accurately quantify the underlying cell-type fractions in a wide range of complex tissues [20-23]. Not until recently, the main limitation has been the availability of a high-resolution tissue-specific DNAm reference matrix, containing representative DNAm profiles for all cell-types in the tissue of interest and which is required by reference-based cell-type deconvolution methods to infer the underlying cell-type fractions [2, 22-27].

Here we use the Illumina 850k DNAm profiles of cell-sorted samples from Salas et al [28] to build a novel 12 immune cell-type DNAm reference matrix for blood tissue, using an improved procedure that exclusively uses cell-type specific unmethylated markers [29]. We validate this 12 immune cell-type DNAm reference matrix on DNAm data with matched flow-cytometric counts, as well as in a large collection of immune cell sorted samples, including whole-genome-bisulfite sequencing (WGBS) samples from the International Human Epigenome Consortium (IHEC) [30]. Importantly, we collate genome-wide DNAm data for a total of 22 independent cohorts, encompassing over 25,000 blood samples, and use our DNAm reference matrix to perform a meta-analysis of immune cell-type fraction associations with common phenotypes, including age, sex, smoking and obesity, validating or strengthening previous findings, whilst also revealing novel associations. In a large cohort with extensive epidemiological and health outcome annotation for approximately 5000 whole blood samples, we identify additional associations of immune-cell fractions with exercise, alcohol consumption, stress and health outcomes, including all-cause mortality. In summary, we use a high quality high-resolution DNAm reference matrix to comprehensively map associations of 12 immune cell-type fractions with common phenotypes and health outcomes.

## Results

### Construction and validation of a 12 blood cell subtype DNAm reference matrix

We constructed a novel DNAm reference matrix encompassing 12 blood cell subtypes (monocytes, neutrophils, eosinophils, basophils, naïve and memory CD4+ T-cells, naïve and memory CD8+ T-cells, naïve and memory B-cells, natural killer (NK) and T-regulatory (Tregs) cells) using the EPIC DNAm profiles of FACS-sorted cells from Salas et al [28]. Since cell-type specific marker genes display a very strong preference for unmethylated promoters and enhancers in the corresponding cell-types [24, 29], we decided to construct a DNAm reference matrix focusing on CpGs specifically unmethylated in each cell-type (**Methods**). For each of the 12 cell-types, we selected CpGs significantly hypomethylated (FDR<0.001) in the given cell-type relative to all the rest, subsequently ranking them according to the difference in average DNAm, so as to ensure maximum separability (**Fig.1a, Methods**). We verified that by selecting the 50 top-ranked CpGs for each cell-type (i.e. a total of 600 CpGs), that these displayed very significant hypomethylation and relatively big differences in average DNAm, as required **(Fig.1a, Additional File 1: table.S1**). About half of the 600 CpGs mapped to gene-bodies, whilst the other half mapped predominantly to inter-genic regions and shores/shelves upstream of the TSS (**Fig.1a**).

**Figure-1:**
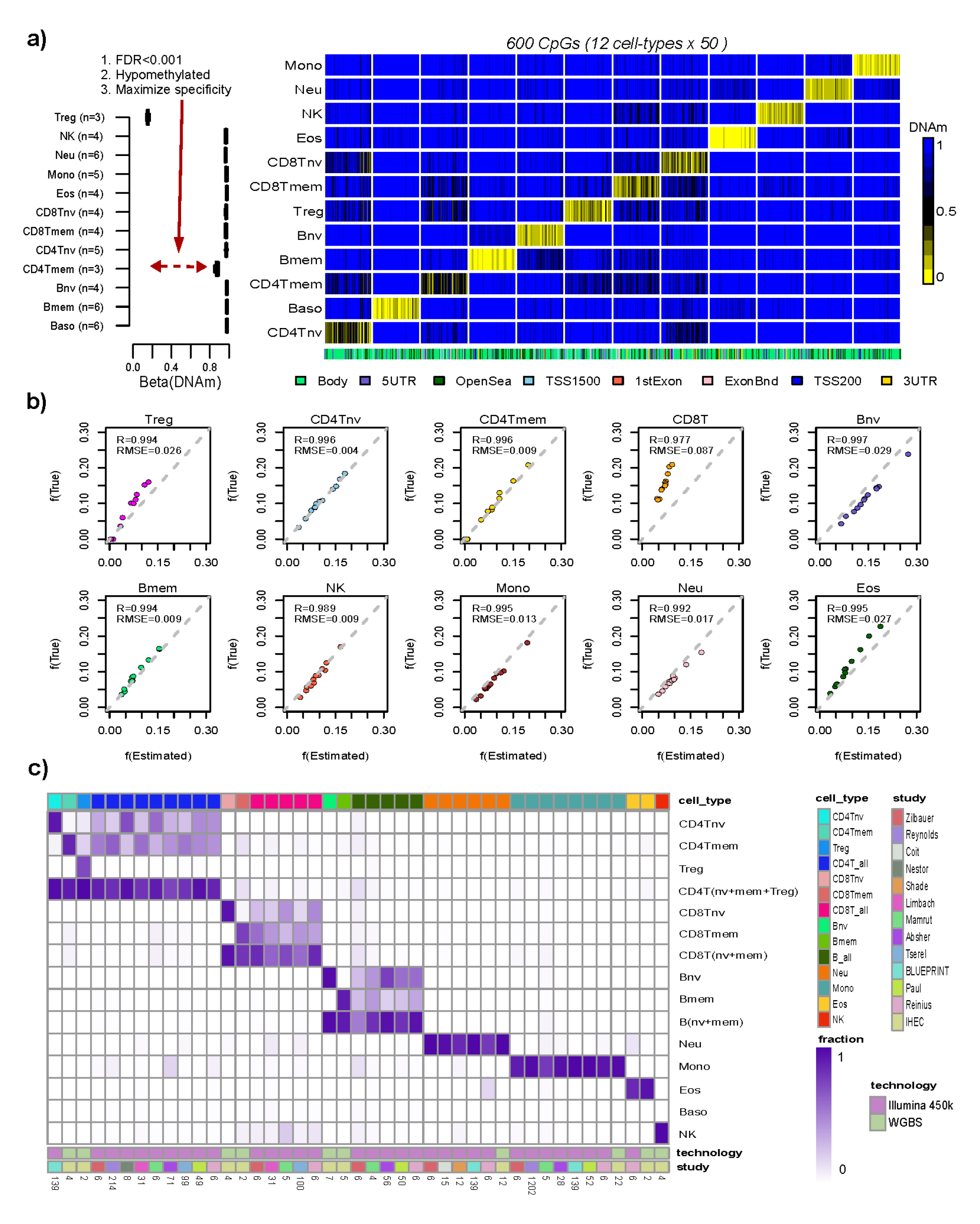
Construction and validation of the 12 blood cell-type hypoDNAm reference matrix. **a) Left panel:** Example of a CpG’s DNAm profile in the DNAm reference matrix, marking T-regulatory (Treg) cells. The y-axis labels the cell-types, x-axis labels the DNAm value, and the number of samples of each cell-type (i.e. in each boxplot) is shown on the y-axis. **Right panel:** The DNAm reference matrix for 12 blood cell subtypes encompassing 600 CpGs (i.e. 50 markers per cell-type). **b)** Scatterplots of true fractions vs estimated fractions for 10 blood cell subtypes using the EPIC DNAm data from 10 artificial mixtures where the underlying mixing proportions were known. For each estimated cell-type we display the R- value (Pearson Correlation Coefficient) and root mean square error (RMSE). **c)** Heatmap displays the estimated fractions of cell-sorted samples for each of the 12 immune cell subtypes in our Illumina 450k/850k DNAm reference matrices, as well as the total CD4+ T-cell, total CD8+ T-cell and total B-cell fractions. The immune cell-type of the sorted sample is indicated by the color-bar on top of the heatmap. The study from which the sorted sample derives from is indicated by the color bar below the heatmap. The technology used to generate the DNAm data of the sorted sample is also indicated. For the 450k and WGBS samples, we used the 450k and 850k DNAm reference matrices, respectively, to obtain the fractions. The estimated fractions in the heatmap are median values taken over biological replicates of the cell-sorted samples, with the number of corresponding biological replicate samples indicated at the bottom.

To validate the 12 blood cell-subtype DNAm reference matrix, we applied it in conjunction with the Robust Partial Correlation (RPC) framework implemented in EpiDISH [21] to estimate cell-type fractions in 12 artificial mixtures where the proportions of 10 underlying blood cell-subtypes used in these mixtures are known [28]. Pearson correlations and root mean square error (RMSE) were excellent for all tested cell-types (**Fig.1b**). To further validate it, we compared estimated cell-type fractions with matched flow-cytometric counts in two independent datasets, one encompassing 6 blood samples with matched counts for 7 blood cell-subtypes [25], and another encompassing 144 peripheral blood samples with matched counts for 3 types of lymphocytes (**Methods**). For both datasets, Pearson correlation R-values and RMSE were reasonably good, further attesting to the quality of our DNAm reference matrix (**Additional File 2: fig.S1a-b**). For completeness, we repeated the construction of a 12 cell-type DNAm reference matrix, but this time restricting to Illumina 450k probes, resulting in a separate 600 CpG x 12 cell-type DNAm reference matrix (**Additional File 1: table S2**). We successfully validated this DNAm reference matrix in artificial blood mixtures and in blood samples with matched flow-cytometric counts (**Additional File 2: fig.S2**). Finally, we further validated both 850k and 450k versions of the DNAm-reference matrix in a large collection of sorted immune-cell subsets, including whole-genome bisulfite sequencing (WGBS) samples from IHEC [31] (**Fig.1c**). Of note, our 850k DNAm reference matrix achieved remarkably high accuracy (mean R-value >0.95) on *in-silico* mixtures generated from these WGBS immune-cell sorted samples (**SI fig.S1c, Methods**), thus demonstrating that our DNAm reference matrix built with Illumina DNAm data is applicable to WGBS data.

### A meta-analysis of immune-cell fractions reveals novel associations with age

We next applied the 12 immune-cell type DNAm reference matrix to perform a large meta-analysis of immune-cell fractions with common phenotypes. This meta-analysis serves two purposes. First, because large DNAm datasets with matched flow-cytometric counts for as many as 12 immune cell-types are not available, it is paramount to seek additional means to validate our high resolution DNAm reference matrix on real data. By estimating immune-cell fractions in a large number of independent cohorts and correlating these to specific phenotypes we can ascertain the quality of our novel DNAm reference matrix. For instance, our DNAm reference matrix should be able to correctly capture a well-known age-associated immuno-senescence signature characterized by a decreased naïve to mature T-cell fraction ratio [6, 32] [33-35] [36, 37]. Second, a meta-analysis can reveal subtle, novel and highly statistically significant associations, not evident from individual studies. To perform the meta-analysis, we estimated fractions for all 12 immune cell subtypes in 22 independent whole blood cohorts, encompassing 25,629 samples and two versions of the Illumina DNAm array (EPIC & 450k) (**Additional File 1: table S3**), and then correlated these fractions to common phenotypes in each cohort using multivariate linear regression models (**Methods**). We used Stouffer’s method to derive an overall z-statistic and P-value of association across all studies with available phenotype information (**Methods**).

We first considered the case of age. The chronological age distribution was reasonably wide for all 22 studies, except for one where all individuals were of the same age and which was henceforth excluded from age-association analyses **(Additional File 2: fig.S3, Additional File 1: table S3**). Validating our DNAm reference matrix, we observed a strong consistent reduction in the naïve CD8+ T-cell population with age across all 21 studies (**Fig.2a**, Stouffer Z=-53, P<10^-200^). For CD4T+ cells, the reduction of the naïve subset was also evident in 17 out of 21 studies (**Fig.2a,** Stouffer Z=-25, P<10^-100^). Correspondingly, there was a trend for the memory T-cell subsets to increase with age (**Fig.2a,** Stouffer Z=14, P<10^-40^ for CD4Tmem and Z=9, P<10^-20^ for CD8Tmem). Of note, effect sizes were small: for instance, in the TZH cohort the CD4T-cell and CD8T-cell naïve fractions displayed reductions of 2% and 1.5% over the course of 50 years (from 20 to 70 years of age).

**Figure-2:**
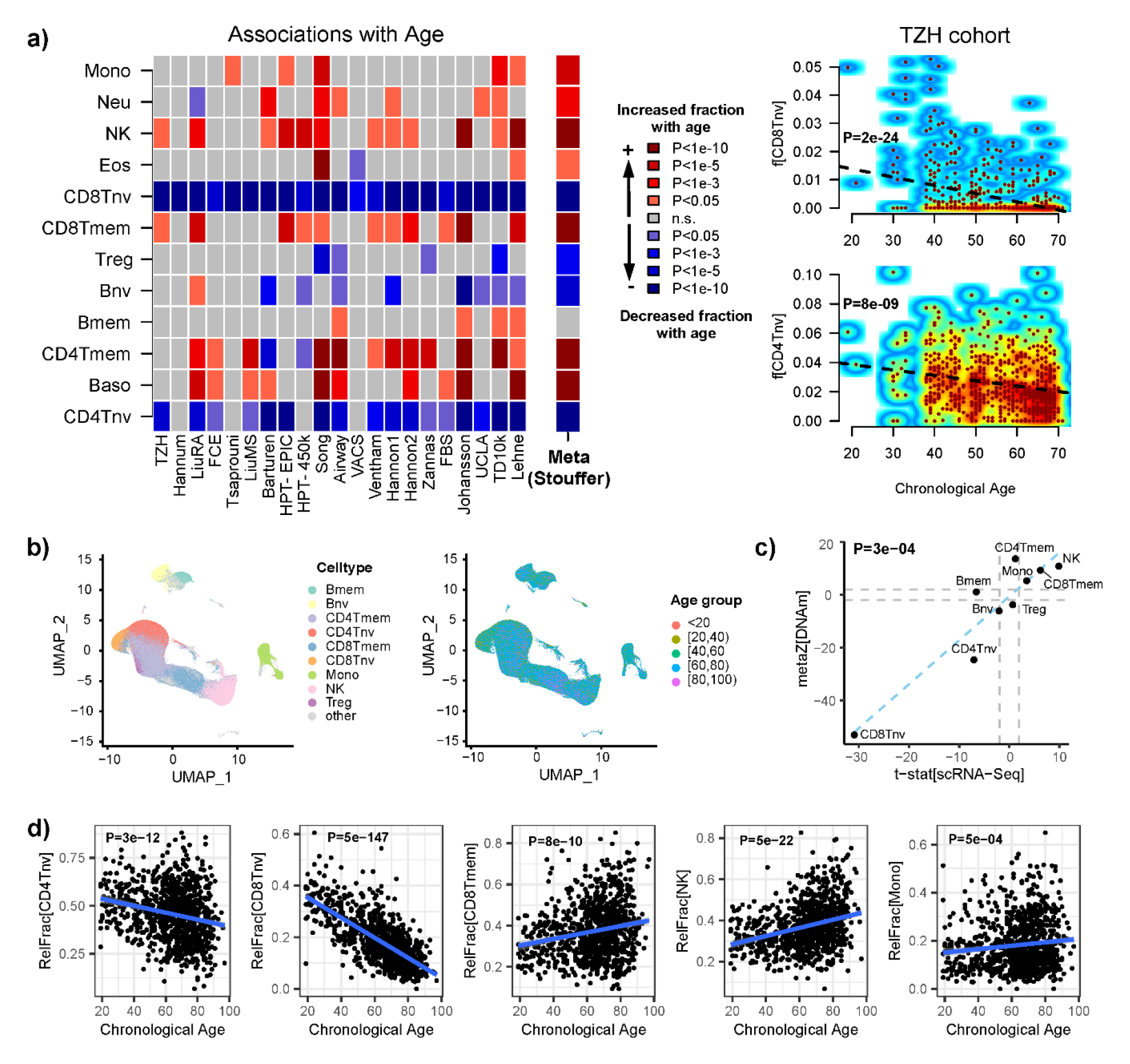
Association and validation of immune cell-type fractions with age. **a)** Heatmap of associations between blood cell-type fractions and age in each of 21 cohorts. Colors indicate both directionality of change and statistical significance, as indicated, with the P-values derived from a multivariate regression that included sex, smoking status and BMI whenever available. Meta(Stouffer) indicates the Stouffer meta-analysis z-statistic and statistical significance. Right panels are smoothed scatterplots displaying the naïve CD8+ and CD4+ T-cell fractions with age in the TZH cohort. Black dashed line and P-value is from a linear regression. **b)** UMAP plots of the scRNA-Seq data of Yazar et al profiling >1 million PBMCs from over 900 donors of different ages. **c)** Scatterplot of Stouffer z-statistics for each cell-type derived from DNAm data, against the corresponding t-statistic from Propeller method based on the scRNA-Seq data. Linear regression P-value of agreement is given. **d)** Examples of immune-cell fractions displaying significant associations with chronological age as inferred from scRNA-Seq data. P-values derive from Propeller.

A meta-analysis over many datasets can also reveal novel associations or strengthen previous preliminary findings. For instance, our meta-analysis revealed a clear trend for basophil and NK cell fractions to increase with age (**Fig.2a,** Stouffer Z=11, P<10^-20^ for NK and Z=10, P<10^-20^ for basophils), strengthening preliminary findings from others [38-40] [41]. To further validate the observed associations for lymphocytes and monocytes, we collated a large scRNA-Seq dataset of over 1.27 million peripheral blood mononuclear cells (PBMCs) from over 900 donors spanning a wide age-range (**Fig.2b**) [42], and used the propeller DA-testing method [43] to derive statistics of association of relative immune-cell fractions with age. This revealed an excellent agreement between the predictions from DNAm data and scRNA-sequencing (**Fig.2c-d**). Our meta-analysis also revealed that the eosinophil fraction did not change significantly with age (**Fig.2a**, Stouffer P=0.03), the marginal significance being driven entirely by one study (Song et al [44]). Of note, this study had profiled DNAm in childhood cancer survivors, and the observed increased eosinophil count in Song et al was independent of cancer-treatment type (**Additional File 2: fig.S4**). Our eosinophil meta-analysis result validates a major eosinophil count study in over 11,000 subjects, which concluded that eosinophil counts do not change with age beyond the age of puberty [45]. Consistent with this, all cohorts analyzed here did not include samples pre-puberty except for the large TD10k cohort which however only included 14 samples younger than 15 years (0.1% of the total) **(Additional File 2: fig.S3b**).

### Decrease of naïve CD8+ T-cell fraction with age is strongest before the age of 40

A large collection of DNAm datasets in blood can also help address the question if age-associated changes in immune-cell fractions is a linear process. We focused on the naïve and memory T-cell fractions because these displayed the most consistent and significant changes with age. To address the question, we merged the estimated fractions of samples in the age range 20 to 85 from all cohorts where the given cell-type fraction was significantly associated with age (with same directionality), excluding the Song et al cohort since this was a PBMC dataset displaying significantly different levels of cell-type fractions (**Additional File 2: fig.S5**). In the case of the naïve CD8+ T-cell fraction, we observed that the decrease was more accentuated for younger individuals in the age-range 25-40, with a shift in the gradient at approximately age-40, followed by a more moderate decrease in the age-range 40 to 60, with no clear further decrease beyond the age of 60 (**Fig.3a**). We verified that this pattern was present in both sexes separately (**Fig.3a**) and in each of the largest cohorts with sufficient and balanced representation across age-groups, demonstrating that the more rapid decrease before age-40 is not a technical artefact of the merging procedure (**Additional File 2: fig.S6**). Correspondingly, the CD8+ T-cell memory fraction displayed a more rapid increase before age-40 (**Fig.3c**). In contrast to the naïve CD8+ T-cell fraction, the naïve CD4+ T-cell fraction displayed a more constant rate of decrease throughout life (**Fig.3b**).

**Figure-3: Non-linear.**
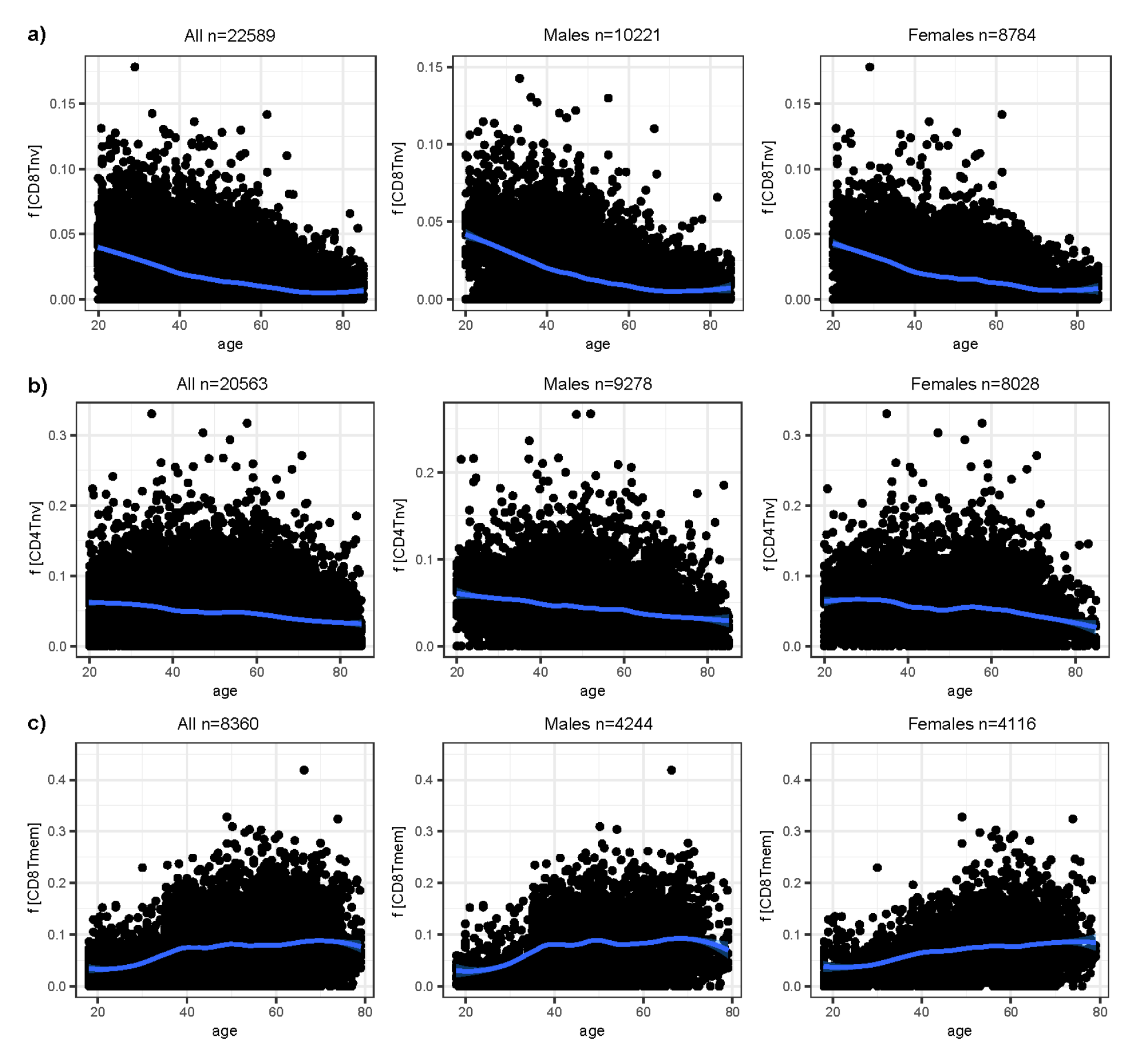
rate of change of immune cell-type fractions with age. **a)** Left panel: Scatterplot of the naïve CD8+ T-cell fraction against chronological age for all samples from all cohorts except Song et al, with the loess regression curve (span=0.3) displayed in blue. Due to large sample size the 95% confidence intervals are generally too tight to be visible. Middle and Right panels: as left but plotting males and females separately. **b)** As a) but for the naïve CD4+ T-cell fraction. Only cohorts where the association of naïve CD4+ fraction with age was significant and with same directionality were used. **c)** As a), but for the CD8+ memory T-cell fraction. Only cohorts where the association of CD8+ memory fraction with age was significant and with same directionality were used.

### Meta-analysis reveals novel associations of immune cell counts with sex

A total of 18 studies profiled blood in both males and females (**Additional File 2: fig.S7a**). The meta-analysis revealed increased monocyte, eosinophil, basophil and NK-cell fractions in males, and increased T-regulatory and naïve T-cell subset fractions in females (**Fig.4a,** Stouffer P<10^-10^ for all cell-types). Once again, we were able to validate most of these associations using the large scRNA-Seq dataset of PBMCs from Yazar et al [42] (**Fig.4b-d**). The observed increased eosinophil fraction in males is also consistent with a similar finding from a major eosinophil count study [45]. Of particular importance is the observed increased regulatory and naïve T-cell fractions in women (**Fig.4a**), an observation that, surprisingly, has not been noted before, except for a sporadic mention in one recent study by Bergstedt et al [46]. Of note, these particular sex-associations were validated using an orthogonal technology (scRNA-Seq data) (**Fig.4c-d**), in clear support of their significance and biological relevance, highlighting a novel insight which could be important for understanding sex-specific differences in cancer and autoimmune disease risk [47]. Interestingly, in the only cohort where the association of naïve CD4+ T-cells with sex was not seen (the Han Chinese TZH cohort, **Fig.4a**), we verified that this was due to residual confounding between alcohol consumption and sex (Chinese women consume little alcohol), as indeed the association became significant when adjusting additionally for alcohol consumption (**Additional File 1: table S4**).

**Figure-4:**
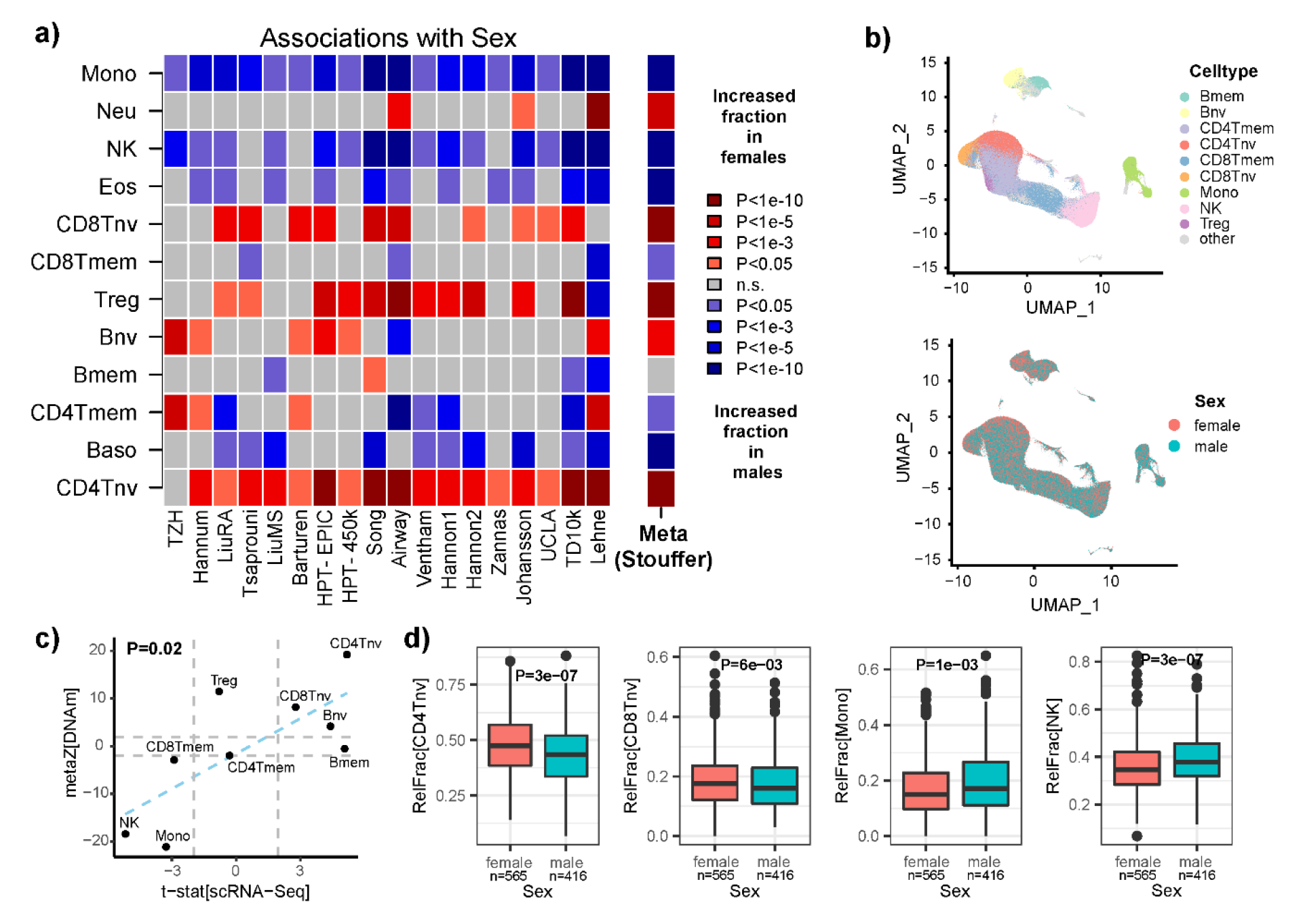
Association and validation of immune-cell fractions with sex. **a)** Heatmap of associations between blood cell-type fractions and sex in each of 18 cohorts with gender information. Red (blue) tones indicate fractions that increase in females (males). P-values derived from a multivariate regression that included age, smoking status and BMI whenever available. Meta(Stouffer) indicates the Stouffer meta-analysis z-statistic and statistical significance. **b)** UMAP plots of the scRNA-Seq data of Yazar et al profiling >1 million PBMCs from over 900 donors encompassing both sexes as shown. **c)** Scatterplot of Stouffer z-statistics for each cell-type derived from DNAm data, against the corresponding t-statistic from Propeller method based on the scRNA-Seq data. Linear regression P-value of agreement is given. **d)** Examples of immune-cell fractions displaying significant associations with sex as inferred from scRNA-Seq data. P-values derive from Propeller.

### Associations of immune-cell fractions with smoking, obesity, exercise and stress

Smoking status information was available for 10 studies (**Additional File 2: fig.S7b**). Compared to age and sex, the number of associations with smoking was much lower (**Fig.5a**). The most significant and consistent associations were displayed by the CD4+ T-cell memory fraction, which was significantly higher in smokers in at least 6 of the 10 studies (**Fig.5a**, Stouffer Z=6, P=4 × 10^−9^). In 5 of 10 studies, we also observed a consistent decrease in the NK-cell fraction in smokers (**Fig.5a**, Stouffer Z=-5, P= 2 × 10^−6^). Although smoking information was not available for the scRNA-Seq study of Yazar et al, so as to validate these results, the findings are nevertheless consistent with a previous blood cell-count study reporting an increased CD4+ T-cell memory fraction in smokers [48] and a report of decreased cytotoxic NK counts in the blood of individuals afflicted with smoking-related diseases like cancer [38]. BMI information was available for 5 studies (**Additional File 2: fig.S7c**), but the only highly significant associations with BMI were seen in the largest of these cohorts (TD10k), encompassing over 6552 samples (**Additional File 2: fig.S8**). For the TD10k cohort additional covariate information was available for most of these 6552 samples, hence for this cohort we performed a more extensive multivariate linear regression analysis including not only age, sex and smoking status as covariates, but also sleep, physical exercise, caffeine and alcohol consumption, and maternal smoking exposure. Interestingly, as with smoking, we observed an increased CD4+ T-cell memory fraction in obese individuals (**Fig.5b,** linear regression P=2× 10^−8^), which was also seen in the second largest cohort with BMI information (TZH cohort) (**Additional File 2: fig.S8**). Of note, in the TD10k cohort, naïve T-cell and NK cell fractions were reduced in obese individuals independently of all other covariates (**Fig.5b,** linear regression P=0.0004 (CD8Tnv), P=0.015 (CD4Tnv) and P=0.0004 (NK)). The multivariate analysis in TD10k also revealed a significant increase of NK (linear regression P=5 × 10^−5^) and monocyte fractions (linear regression P=0.0009) in individuals undergoing frequent exercise, whilst the naïve B-cell fraction decreased (linear regression P=0.0001) (**Fig.5b**). The NK-fraction also increased in individuals sleeping longer hours although this association was more marginal (linear regression P=0.003, **Fig.5b**). In contrast, the NK fraction decreased significantly in individuals reporting higher stress levels (linear regression P=0.0006, **Fig.5b**). Although associations with alcohol consumption were only of marginal significance, it is worth noting that the specific increases of CD4T cells (both naïve and memory subsets) and basophils in heavy drinkers was replicated at a similar significance level in the TZH cohort (**Additional File 2: fig.S9**). Overall, these patterns reveal associations of specific cell-types (e.g. memory and naïve T-cells, NK) with multiple independent factors (smoking, obesity, stress, exercise and spirit drinking).

**Figure-5:**
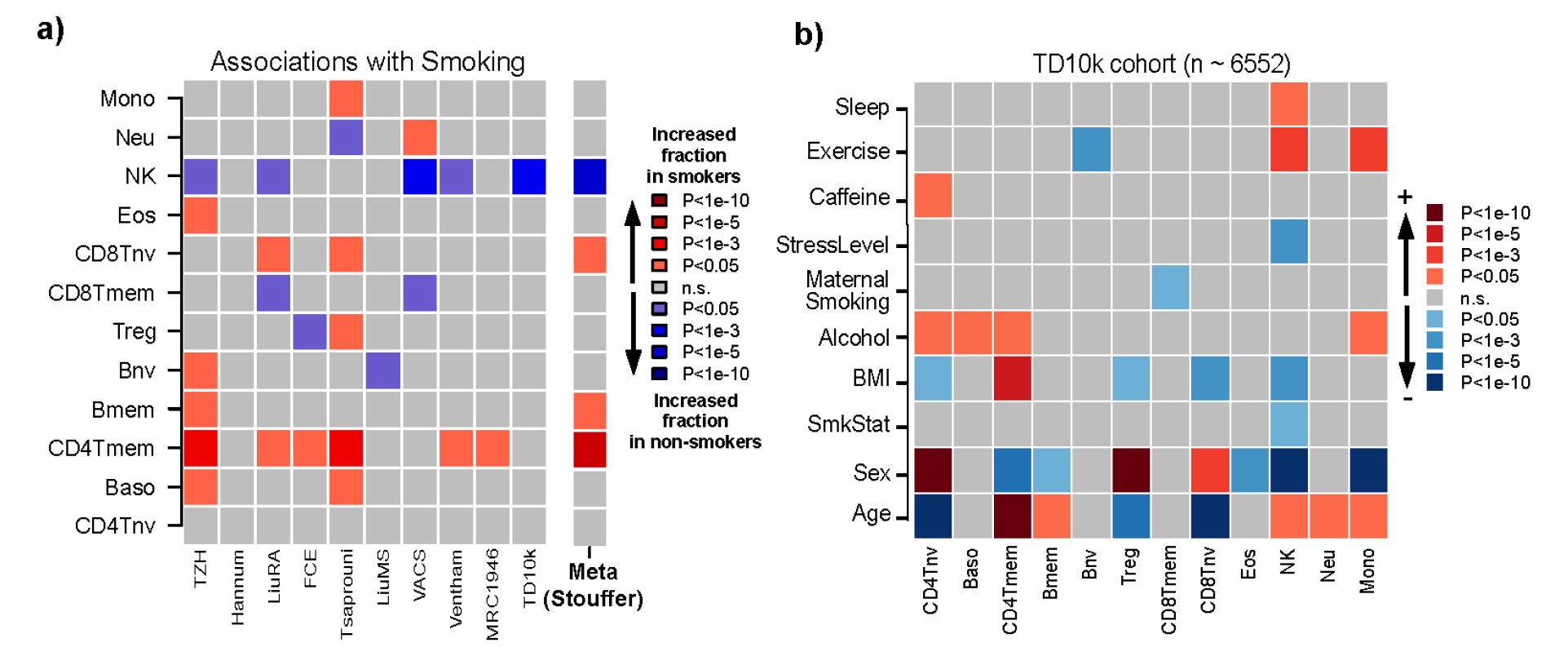
Association and validation of immune-cell fractions with smoking, BMI, exercise and stress. **a)** Heatmap of associations between blood cell-type fractions and smoking in each of 10 cohorts with smoking-status information. Red (blue) tones indicate fractions that increase in smokers (never-smokers). P-values derived from a multivariate regression that included age, sex and BMI whenever available. Meta(Stouffer) indicates Stouffer z-statistic and significance. **b)** Heatmap of multivariate associations of immune-cell fractions with epidemiological factors in the TD10k cohort, encompassing over 6552 samples. Red tones indicate fractions that increase with increasing values of the epidemiological factors, blue tones indicate decreases. P-values were derived from a multivariate linear regression including all phenotypic factors as shown.

### Associations of immune-cell fractions with health outcomes

In order to explore associations with health outcomes, we focused on a subset of the TD10k cohort derived from the Mass General Brigham Biobank (**Methods**). This subset has extensively annotated epidemiological and prospective health outcome information, including all-cause mortality, type-2 diabetes (T2D), cancer, chronic obstructive pulmonary disease (COPD), stroke, cardiovascular disease (CVD) and depression (**Methods**) for 4386 subjects who were healthy at blood sample draw. The demographics of this cohort is shown in **Additional File 1: table S5**. In the first instance, we used Cox-proportional hazard regressions to evaluate associations between each of the 12 immune-cell fractions with each of the various outcomes, adjusting for age, sex and race (**Methods**). We observed many associations (**Fig.6a, Additional File 1: tables S6-S7**) that remained significant upon further adjustment for additional disease risk factors including smoking, obesity and alcohol consumption (**Fig.6b, Additional File 1: tables S8-S9**). Notably, naïve CD4+ T-cell, naïve B-cell and NK-cell fractions were all associated with a reduced risk of all-cause mortality, even after adjustment for all major disease risk factors. Interestingly, whilst the naïve CD4+ T-cell fraction also displayed negative associations with many health outcomes, notably with COPD and T2D, the memory CD4+ T-cell fraction was only negatively associated with all-cause mortality. Some of the other associations were strongly dependent on the specific health outcome. For instance, an increased memory B-cell fraction was specifically associated with an increased risk of cancer but a reduced risk for T2D, whilst no associations were observed for the other outcomes. These results highlight the importance and value of immune-cell fractions as biomarkers of disease risk.

**Figure-6:**
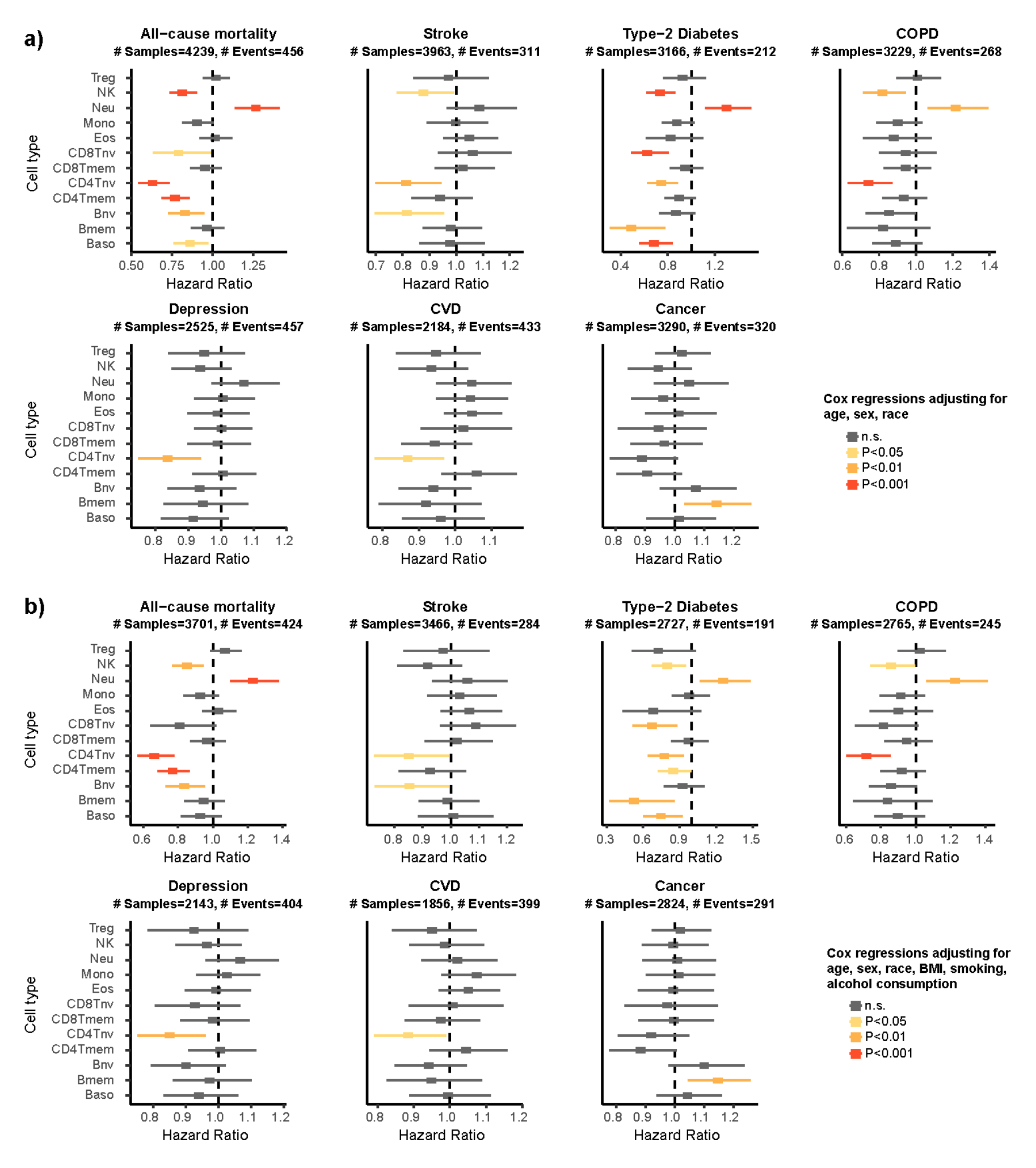
Association of immune-cell fractions with health outcomes in TD10k cohort. **a)** Forest plots of association between immune-cell fractions with various health outcomes, as shown. In each panel, the x-axis labels the Hazard Ratio (HR) as evaluated using a Cox-regression model that included age, sex and race as covariates. P-values derive from the Wald-test. Vertical dashed line indicates the line HR=1. For each HR datapoint, we display the 95% confidence interval. For each health outcome we display the total number of samples and events. **b)** As a) but adjusting in the Cox-regression model also for smoking, BMI and alcohol consumption.

Next we asked how well we would be able to predict all-cause mortality using a model that includes all immune-cell fractions in addition to age, gender, race, smoking, BMI and alcohol consumption. To this end, we split the dataset into a 70% training (2591samples, 302 events) and 30% test set (1110 samples, 122 events) and trained a lasso Cox-proportional hazards regression model on the training set using a 10-fold internal cross-validation strategy (**Methods**). Using the C-index as the performance metric we identified two optimal models with overlapping 95% confidence intervals: one included only age and naïve CD4+ T-cell fraction as covariates, and another included all variables excluding neutrophil and CD8+ T-cell fractions (**Additional File 1: table S10**). These models achieved Hazard Ratio (HR) and C-index values of HR=2.99 (95%CI: 2.57-3.48) & C=0.75 (95%CI: 0.72-0.78) and HR=3.58 (95%CI: 3.08-4.16) & C=0.79 (0.77-0.81), respectively (**Additional File 1: table S11**). Evaluation of these two models in the 30% blind test set revealed similar HR and C-index values of HR=3.58 (95%CI: 2.79-4.60) & C=0.77 (95%CI: 0.73-0.81) and HR=3.90 (95%CI: 3.07-4.96) & C=0.80 (95%CI: 0.77-0.84), respectively (**Additional File 1: table S11**). As a benchmark, chronological age alone achieved C-index values of 0.73 (0.71-0.76) and 0.75 (0.71-0.80) in training and test sets, respectively (**Additional File 1: table S11**), indicating that although the association with all-cause mortality was dominated by age, most immune-cell fractions and in particular the naïve CD4+ T-cell fraction, did contribute to an improved predictive model.

### Decreased memory T-cell fractions in severe Covid-19 patients

Finally, we explored associations of immune-cell fractions with Covid-19 disease severity. Barturen et al profiled DNAm in whole blood from 574 Covid-cases (positive at blood sample draw) and controls (negative at blood sample draw) [49]. Multivariate regression analysis adjusting for age and sex revealed multiple associations, with CD4 and CD8 memory T-cells, NK-cells, monocytes and eosinophils all displaying reduced fractions with disease severity, whilst the neutrophil fraction increased (**Additional File 2: fig.S10a**). Whilst the increased neutrophil fraction in severe Covid-19 cases is well-known [49-52], the specific reduction in memory T-cell subsets was not previously noted by Barturen et al. We thus sought independent validation using a scRNA-Seq dataset that profiled immune-cells from severe and mild Covid-19 cases as well as healthy controls (**Methods**) [53]. Once again we used the propeller method [43] to detect differential abundance of immune cell subsets between severe and mild/healthy cases (**Additional File 1: table S12**). This confirmed the decrease in the T-cell memory fraction in severe cases (**Additional File 2: fig.S10b-d**). Of note, an increased T-cell memory fraction is a hallmark of recovery from severe Covid-19 disease [52]. Thus, these data demonstrate the ability of our DNAm reference matrix to find subtle immune-cell shifts with Covid-19 severity.

## Discussion

This work contributes a novel DNAm reference matrix defined over 12 immune cell-types, which is valid for both Illumina and whole-genome-bisulfite sequencing DNAm data. Using this DNAm reference matrix we performed a large meta-analysis of cell-type fractions in blood, in order to comprehensively map associations between these 12 immune-cell fractions and common phenotypes. The meta-analysis served two purposes. First, as a means of further validating our DNAm reference matrix by testing for known associations between immune-cell fractions and a broad range of phenotypes. For instance, further validating the derived fractions for naïve T-cell subsets, we confirmed the reduced naïve to mature CD8+ and CD4+ T-cell ratio with age [6, 32] [33-35]. The increased neutrophil fraction in severe Covid-19 patients is another well-known observation [49-52], which we correctly retrieved, thus further validating the neutrophil component of our DNAm reference matrix. Our meta-analysis also established that the eosinophil count does not change with age but that it is higher in males compared with females, consistent with a previous large eosinophil count study [45]. The higher NK-cell fraction in males compared to females is validated by several reports [50, 65] and a study that used single-cell RNA-sequencing to compare immune-cell fractions between males and females [66]. There is also strong prior evidence for an increased NK-fraction with age [38, 39] [40], and in individuals undergoing frequent physical exercise [54, 55], further validating the NK-component of our reference matrix. The increased monocyte fraction in males compared to females has also been previously observed [56]. In the case of age, sex and Covid-19 disease severity, associations inferred from the DNAm data were strongly validated using independent scRNA-Seq data. Of note, although meta-analysis P-values were highly significant, for certain cell-types individual cohorts displayed differences, likely caused by the small effect sizes involved or to intrinsic differences between studies (e.g sample size), thus highlighting the importance of performing meta-analyses to increase power.

The meta-analysis also revealed a number of biologically and clinically significant insights and connections, which were not previously known, or for which prior evidence was scarce or controversial. For instance, there was little prior evidence for an increased naïve CD4+ and T-regulatory cell fractions in women compared to men, except for one study by Bergstedt et al [46]. A recent review reported contradictory findings for regulatory T-cells in mice [50], and two small-sized studies reported increased T-regulatory counts [65] and naïve T-cell fractions [38] in males. In light of this controversy, our meta-analysis serves to unequivocally demonstrate that the naïve and regulatory T-cell fractions are higher in women, a result that we further validated using independent scRNA-Sequencing data. Of note, in females naïve CD4+ T-cells preferentially produce IFNγ upon stimulation, whereas in males they produce more IL- 17 [57]. Correspondingly, IFNγ production has been found to be higher in females [58]. Given IFNγ’s major role in activating anticancer immunity, it is therefore plausible that this subtle increase in the naïve CD4+ T-cell fraction could contribute to the well-known overall lower cancer-incidence in women [59].

Another insight, which was however only seen in the largest cohorts with available BMI information, is the increased memory CD4+ T-cell fraction in obese individuals, consistent with a previous study [60] and another reporting an increased circulating CD4+ T cell frequency with increased BMI [61]. The increased CD4+ T-cell memory fraction in obese individuals could reflect increased activation due to adipocyte antigen presenting cells [61] and inflammation [62] within adipose tissue. Another interesting connection was centered on the NK-fraction, a key component of anti-cancer immunity, which displayed decreases with cancer risk factors such as smoking, obesity and stress levels, whilst it increased with cancer-preventive factors such as exercise and hours of sleep, all these associations being derived from multivariate models that included age and sex. On the other hand, the NK-fraction increased with age and was higher in males compared to females. These data clearly indicate the importance of recording all epidemiological factors in individual studies, as multiple factors can impinge on measured cell-type fractions.

Another noteworthy insight was the observation of an increased eosinophil count with age in the blood of childhood cancer survivors, when this increase was not evident in any of the other 20 cohorts. This observation could be significant as childhood cancer survivors are known to be at a much higher risk of developing heart disease [44, 63], and one particular rare condition that can lead to a range of cardiovascular disease manifestations is eosinophilic myocarditis [64], a condition associated with elevated eosinophil counts. Thus, our finding leads to the hypothesis that the increased age-associated cardiovascular disease risk in childhood cancer survivors could be driven in part by an age-associated increase in eosinophils. What triggers this increase in the eosinophil fraction with age is unclear however, as the association was independent of the type of cancer treatment received (i.e. chemo or radiotherapy). However, we also express caution because the childhood cancer study profiled PBMCs, which is depleted for granulocytes, and so the observed association between eosinophil counts and age could also be due to differences in mononuclear cell isolation efficiency.

By merging cell-type fractions from different cohorts together, we were also able to establish that the naïve CD8+ T-cell fraction decreases non-linearly with age, displaying a more pronounced decrease in the age-range 20 to 40 compared to mid-life (40-60 years), with the decrease being much less noticeable beyond 65 years of age. In contrast, the naïve CD4+ T- cell fraction displayed a linear decrease throughout life. Elucidating the biological basis of these differences could have important ramifications for our understanding of the aging immune-system. Interestingly, of all immune-cell fractions, the naïve CD4+ T-cell fraction displayed the strongest association with all-cause mortality, with increased fractions associated with a significantly reduced risk, which we note was independent of all major disease risk factors including age, sex, race, smoking, BMI and alcohol consumption. Correspondingly, the naïve CD4+ T-cell fraction also displayed significant associations with individual health outcomes including COPD, CVD and T2D. Likewise, the NK-fraction was associated with a reduced risk of all-cause mortality as well as COPD and T2D. Considering individual health outcomes was important: for instance, whilst the memory B-cell fraction did not correlate with all-cause mortality, an increase in this fraction was associated with an increased risk of cancer and simultaneously with a reduced risk of T2D. In relation to cancer-risk, the increased memory B-cell fraction could be associated with reduced anti-tumor immunity through increased release of pro-tumorigenic factors such as IL-10 or TGF-beta [65].

## Conclusions

In summary, this study has comprehensively mapped associations between immune-cell fractions and common phenotypes at an unprecedented high resolution of 12 immune cell subtypes in blood, revealing many important associations with factors such as age, sex, smoking, obesity and health outcomes. The DNAm reference matrix encompassing 12 immune-cell subtypes that we present here has been extensively validated, including WGBS data, and is made freely available as part of our EpiDISH Bioconductor R-package. We envisage that similar high resolution meta-analyses performed in tissues other than blood using tools such as EpiSCORE [22, 24] could help discern changes in cell-type composition that are important predictors or contributors of disease and disease risk.

## Methods

### Construction of the 12 immune cell-type DNAm reference matrices

We obtained the EPIC DNAm dataset of 12 sorted immune-cell subsets from GEO under accession number GSE167988. The idat files were downloaded and processed using *minfi* R-package [66]. We retained 756625 probes with significantly detected values across all 68 samples. The resulting beta-valued data matrix was then adjusted for type-2 probe bias using BMIQ [67]. We next removed 12 artificial mixture samples, leaving a total of 56 sorted samples: 6 basophils (Baso), 6 memory B-cells (Bmem), 4 naïve B-cells (Bnv), 4 memory CD4+ T-cells (CD4Tmem), 5 naïve CD4+ T-cells (CD4Tnv), 4 memory CD8+ T-cells (CD8Tmem), 5 naïve CD8+ T-cells (CD8Tnv), 4 eosinophils (Eos), 5 monocytes (Mono), 6 neutrophils (Neu), 4 natural killer (NK) and 3 regulatory T-cells (Treg). We then performed SVD, estimating the number of significant components using RMT [68], followed by hierarchical clustering on the sample projection matrix, to check that sorted samples clustered by cell-type. This revealed that one CD4Tmem and one CD8Tnv sample did not cluster correctly. Hence, these two samples were removed leaving a normalized beta-valued data matrix over 756,625 probes and 54 sorted samples. We then used *limma* [69, 70] to perform differential DNAm analysis for each of the 12 cell-types in turn, comparing that cell-type to the other eleven. Next, for each cell-type we selected those probes passing a Benjamini-Hochberg adjusted FDR<0.001 and displaying hypomethylation in the given cell-type compared to the rest. We only consider hypomethylated probes because these are overwhelmingly more likely to be truly cell-type specific markers [24, 29]. The above procedure can still result in probes attaining smaller DNAm-values in another cell-type because the limma analysis compares an average of the given cell-type to the average over 11 other cell-types. Hence, to ensure that our selected probes for a given cell-type attain the smallest DNAm values in that cell-type, we also recorded for each cell-type *t* and probe *p* the minimum DNAm difference value (called Δ_*pt*_) between the cell-type of interest *t* and the other 11 cell-types. We note that for the probes of interest these values will be negative because the maximum value across the samples of the given cell-type of interest should be lower than the minimum value across all other cell-types. Then for all hypomethylated probes at FDR<0.001 for a given cell-type *t*, we ranked these in increasing order of the Δ_*pt*_ values, to ensure that the top-ranked probes display the largest negative Δ_*pt*_ values. This means that for these probes the maximum beta DNAm-value across the samples of the given cell-type of interest is much smaller than the minimum value across the samples from all other cell-types, which ensures that we are selecting probes with the largest effect sizes. For each cell-type we then selected the top-ranked 50 probes as cell-type specific markers, resulting in a total of 600 (50 times 12) unique marker probes. We note that the number of hypomethylated probes at FDR<0.001 per cell-type was in general quite large: mean over the 12 cell-types was 24954, range was 1111 (CD4Tmem) to 88992 (Bmem). Hence, selecting the top-ranked 50 according to Δ*_pt_* values ensures that we are selecting not only highly significant hypomethylated probes but also those with the largest possible effect sizes. The final DNAm reference matrix over the 600 marker probes was then built by taking the median DNAm value over the samples of a given cell-type. Note that we take the median, because this is a more robust estimator and because later we estimate cell-type fractions using a robust partial correlation framework which does not require the assumption that the reference value should be an average (in contrast to constrained projection which does). In order not to bias performance in Illumina 450k datasets, we also generated a separate 12 cell-type DNAm reference using only 450k probes, using the exact same procedure as described above. Of note this 450k DNAm reference matrix is also defined for 600 unique marker probes.

### Validation DNA methylation datasets of sorted samples

We obtained independent immune cell sorted samples from the following sources, encompassing both Illumina 450k and WGBS technologies: From Reynolds et al [71] we obtained 1202 monocyte and 214 CD4+ T-cell 450k samples (GEO: GSE56581). From BLUEPRINT [72] we obtained 139 monocyte, 139 naïve CD4+ T-cell and 139 neutrophil 450k samples from the same 139 individuals. From Zilbauer et al we obtained 6 CD4+ T-cell, 6 CD8+ T-cell, 6 B cell, 6 neutrophil and 6 monocyte 450k samples (ArrayExpress: E-MTAB- 2145). From Coit et al [73] we obtained 15 neutrophil 450k samples (GEO: GSE65097). From Nestor et al [74]we obtained 8 CD4+ T-cell 450k samples (GEO: GSE50222). From Shade et al [75]we obtained 12 neutrophil 450k samples (GEO: GSE63499). From Limbach et al [76] we obtained 31 CD4+ T-cell and 31 CD8+ T-cell 450k samples (GEO: GSE71955). From Mamrut et al [77] we obtained 6 CD4+ T-cell, 5 CD8+ T-cell, 4 B cell and 5 monocyte 450k samples (GEO: GSE71244). From Absher et al [78]we obtained 71 CD4+ T-cell, 56 B cell and 28 monocyte 450k samples (GEO: GSE59250). From Tserel et al [79]we obtained 99 CD4+ T-cell and 100 CD8+ T-cell 450k samples (GEO: GSE59065). From Paul et al [80] we obtained 49 CD4+ T-cell, 50 B cell and 52 monocyte 450k samples (EGA: EGAS00001001598). From Reinius et al [81] we obtained 6 CD4+ T-cell, 6 CD8+ T-cell, 6 B cell, 6 neutrophil, 6 monocyte and 6 eosinophil 450k samples. From the IHEC data portal (https://epigenomesportal.ca/ihec/), we obtained WGBS hg38 immune-cell sorted samples from build version 2020-10. The downloaded files were in bigwig format. For each sample, there are 2 bigwig files, one for read coverage information and the other for beta value information. We first used bigWigToWig shell script provided by UCSC genome browser to convert them into wig files. Then for each sample, we combined the read coverage and beta value information into one file and stored them as an .rda file for further processing. For each WGBS sample, we found the CpGs present in the 850k DNAm beadarray. We dropped one sample (ERS568736) due to ultra-low coverage. For the rest of samples, the minimum number of 850k probes covered by 20 reads or more was 336,812, the maximum was 834,149, with a mean of 689,451. In total, for this work we used 4 memory CD4+ T-cell, 2 T-regulatory, 4 naïve CD8+ T-cell, 2 memory CD8+ T-cell, 7 naïve B cell, 5 memory B cell, 12 neutrophil, 22 monocyte, 2 eosinophil and 4 natural killer cell WGBS samples, for validating our 850 DNAm reference matrix.

### In-silico mixture validation analysis with WGBS cell-sorted samples

The overall coverage of common 850k probes across all WGBS samples was only 132,713, containing only 72 probes from our 600 CpG 850k DNAm reference matrix. Hence, for the in-silico mixture analysis we did not impose any threshold on read coverage, which resulted in 487,795 probes, including 304 probes from our 850k DNAm reference matrix. This is sensible because reference-based cell-type deconvolution can tolerate even up to 30% errors in the DNAm reference matrix [82]. Hence, in-silico mixtures were generated from the 4 memory CD4+ T-cell, 2 T-regulatory, 4 naïve CD8+ T-cell, 2 memory CD8+ T-cell, 7 naïve B cell, 5 memory B cell, 12 neutrophil, 22 monocyte, 2 eosinophil and 4 natural killer cell WGBS samples, defined over the 304 probes. We generated 1000 in-silico mixtures, randomly selecting one sample from each immune-cell type and using random weights drawn from a uniform distribution to generate the linear combination. Since there are a total of 4*2*4*2*7*5*12*22*2*4=4,730,880 potential combinations, generating 1000 in-silico mixtures is sensible as the number is large enough to reliably assess performance, whilst also reducing statistical dependency of the combinations as much as possible. Performance was assessed using Pearson R-values and RMSE.

### Illumina DNA methylation datasets used in the meta-analysis

Below we provide details of the data source and processing of each dataset used in our meta-analyses. Each dataset profiled whole or peripheral blood samples with Illumina DNAm beadarrays (EPIC or 450k). Further details are available in **Additional File 1: table S3**.

*LiuMS:* The 450k dataset from Kular et al [83] was obtained from the NCBI GEO website under the accession number GSE106648 (https://www.ncbi.nlm.nih.gov/geo/query/acc.cgi?acc=GSE106648). We downloaded the series matrix file which contained the data processed with detection P-values. We only retained probes with no missing values across all samples. This data was subsequently normalized with BMIQ [67], resulting in a normalized data matrix for 483,567 probes and 279 peripheral blood samples (140 multiple-sclerosis patients + 139 controls).

*Song:* The EPIC dataset from Song et al [44] profiled DNAm in blood from childhood cancer survivors and was obtained from the NCBI GEO website under the accession number GSE169156 (https://www.ncbi.nlm.nih.gov/geo/query/acc.cgi?acc=GSE169156). The file “GSE169156_RAW.tar” which contains the IDAT files was downloaded and processed with *minfi* package [66]. Probes with P values < 0.05 across all samples were retained. The filtered data was subsequently normalized with BMIQ, resulting in a normalized data matrix for 823,395 probes and 2052 samples.

*HPT-EPIC & HPT-450k:* These datasets derived DNAm profiles from the peripheral blood of African-Americans as part of The Genetic Epidemiology Network of Arteriopathy (GENOA) study. Data was obtained from the NCBI GEO websites under the accession numbers GSE210255 (https://www.ncbi.nlm.nih.gov/geo/query/acc.cgi?acc=GSE210255, Infinium HumanMethylationEPIC BeadChip) and GSE210254 (https://www.ncbi.nlm.nih.gov/geo/query/acc.cgi?acc=GSE210254, Infinium HumanMethylation450k BeadChip). The files “GSE210255_RAW.tar” and “GSE210254_RAW.tar” containing the IDAT files were downloaded and processed with minfi package. Probes with P values < 0.05 across all samples were retained. The filtered data was subsequently normalized with BMIQ, resulting in normalized data matrices containing 826,512 probes and 1394 samples (EPIC set), and 476,722 probes and 418 samples (450k set), respectively.

*Barturen:* The EPIC dataset from Barturen et al [49] profiled DNAm in blood from Covid-19 patients with three different levels of disease severity. Data was obtained from the NCBI GEO website under accession number GSE179325 (https://www.ncbi.nlm.nih.gov/geo/query/acc.cgi?acc=GSE179325). The file “GSE179325_RAW.tar” containing IDAT files was downloaded and processed with minfi package. Only probes with P values < 0.05 across all samples were retained. The filtered data was subsequently normalized with BMIQ, resulting in a normalized data matrix for 845,921 probes and 574 samples.

*Airwave*: The EPIC dataset from the Airwave study [84] was obtained from the NCBI GEO website under accession number GSE147740 (https://www.ncbi.nlm.nih.gov/geo/query/acc.cgi?acc=GSE147740). The file “GSE147740_RAW.tar” containing IDAT files was downloaded and processed with minfi. Only probes with P-values < 0.05 across all samples were retained. Filtered data was subsequently normalized with BMIQ, resulting in a normalized data matrix for 840,034 probes and 1129 samples.

*VACS:* The 450k dataset from Zhang X et al [85] was obtained from the NCBI GEO website under accession number GSE117860 (https://www.ncbi.nlm.nih.gov/geo/query/acc.cgi?acc=GSE117860). The file “GSE117860_MethylatedSignal.txt.gz” containing the unmethylated signals, methylated signals and detection P-values was downloaded. The beta values were obtained using the Illumina definition: methylated signal / (methylated signal + unmethylated signal + 100). Only probes with P-values < 0.05 across all samples were retained. The filtered beta value matrix was subsequently normalized with BMIQ, resulting in a normalized data matrix 396,327 probes across 529 samples.

*Ventham:* The 450k dataset from Ventham et al [86] was obtained from NCBI GEO website under accession number GSE87648 (https://www.ncbi.nlm.nih.gov/geo/query/acc.cgi?acc=GSE87648). The file “GSE87648_RAW.tar” containing the IDAT files was downloaded and processed with minfi package. Two samples in which the proportion of probes with P-values < 0.05 is lower than 0.99 were excluded, and probes with P-values < 0.05 across all remaining samples were kept. Filtered data was subsequently normalized with BMIQ, resulting in a normalized data matrix for 470,807 probes and 382 samples.

*Hannon-1 and 2:* The 450k datasets from Hannon et al [87, 88] were obtained from NCBI GEO websites under accession numbers GSE80417 (https://www.ncbi.nlm.nih.gov/geo/query/acc.cgi?acc=GSE80417) and GSE84727 (https://www.ncbi.nlm.nih.gov/geo/query/acc.cgi?acc=GSE84727), which represent the phase-1 and phase-2 of their study. For the phase-1 dataset, the file “GSE80417_rawBetas.csv.gz” containing the beta values for filtered probes was downloaded, and the beta values were then normalized with BMIQ algorithm. For the phase-2 dataset, the file “GSE84727_rawBetas.csv.gz” and “GSE84727_detectionP.csv.gz” were downloaded. Only probes with P-values < 0.05 across all samples were kept. The filtered beta value data was subsequently normalized with BMIQ. The final data matrices contain 477,818 probes across 675 samples, and 478,630 probes across 847 samples, for phase-1 and phase-2, respectively.

*Zannas:* This 450k dataset is derived from whole blood of African American participants of the Grady Trauma Project [89]. Data was obtained from NCBI GEO website under accession number GSE72680 (https://www.ncbi.nlm.nih.gov/geo/query/acc.cgi?acc=GSE72680). The file “GSE72680_beta_values.txt.gz” containing the beta values and detection P-values was downloaded. Probes with P-values < 0.05 across all samples were kept. However, the beta-value matrix of the retained probes still contained NAs and these were imputed with the function impute.knn (k=5) from the *impute* R-package. Beta values were later normalized with BMIQ, resulting in a normalized data matrix containing 453,310 probes across 422 samples. *Flanagan/FBS:* The 450k dataset Flanagan et al is from the Breakthrough Generations Study [90] and was obtained from NCBI GEO website under the accession number GSE61151 (https://www.ncbi.nlm.nih.gov/geo/query/acc.cgi?acc=GSE61151). The file “GSE61151_Matrix_raw_signal.txt.gz” containing beta values and detection P-values was downloaded. Only probes with P-values < 0.05 and with no other QC-failures across all samples were kept. The filtered beta value data was subsequently normalized with BMIQ. The 2 duplicates of 4 pairs of samples were averaged, since duplicate pairs exhibited strongest correlations with each other. The final normalized data matrix was defined for 426,430 probes and 184 samples.

*Johansson:* The 450k dataset from Johansson et al [91] was obtained from NCBI GEO website under accession number GSE87571 (https://www.ncbi.nlm.nih.gov/geo/query/acc.cgi?acc= GSE87571). The file “GSE87571_RAW.tar” containing the IDAT files was downloaded and processed with *minfi* R-package. Probes with P-values < 0.05 across all samples were kept. Filtered data was subsequently normalized with BMIQ, resulting in a normalized data matrix for 475,069 probes across 732 samples.

*TD10k:* This EPIC DNAm dataset from TruDiagnostic Inc. was collected between 2020 – 2022. The dataset represents whole blood samples collected from a total of 10,420 individuals who had provided blood as part of either a routine check by physicians or by acquiring a kit directly from TruDiagnostic Inc (N = 7,023). Another subset of samples (N = 3,397) were retrieved from the Mass General Biobank and sent to TruDiagnostic Inc. All individuals have provided consent to use the collected data for this project. Whole blood samples were collected and stored at − 80C prior to DNA processing, which was conducted at the TruDiagnostic Inc. lab facility (Lexington, KY, USA). 500ng of DNA was extracted and bisulfite converted using the EZ DNA Methylation kit (Zymo Research) using manufacturer’s instruction. After bisulfite conversion, converted DNA were hybridized to the Illumina HumanMethylation EPIC Beadchip, stained, washed, and imaged with the Illumina iScan SQ instrument to obtain raw image intensities. Raw data was processed using the *minfi* pipeline. Low quality samples were identified using the ENmix qcfilter() function. Probes with P-values < 0.05 across all samples were identified and kept, with low quality probesets removed. A combinatorial normalization processing using the minfi Funnorm procedure, followed by the RCP method available in ENmix(). The final normalized beta-valued matrix was defined for 864,627 probes across 10,296 samples.

*Lehne:* This 450k DNAm dataset consists of over 2700 peripheral blood samples [92], but we used the already QC-processed and normalized version previously described by Voisin et al [93] which included a total of 2639 samples.

*UCLA:* The UCLA dataset (N=178) was collected at PhysioAge LLC and sent to TruDiagnostic Inc. for processing. All processing and data normalization was performed exactly as for the TD10k dataset.

*TZH, Hannum, LiuRA, Tsaprouni*, *MRC1946* and *FCE*: The TZH (EPIC), Hannum (450k), LiuRA (450k), Tsaprouni (450k), MRC1946 (450k) and FCE (EPIC) datasets were downloaded and normalized as described by us previously [18, 94, 95].

### Mass General Biobank data (MGB)

In addition to the 3,397 samples that were included in the TD10k cohort, additional samples were retrieved from the MGB Biobank to increase power in the association studies with health outcomes (not part of the meta-analysis), to a total of 4,386 samples. MGB derived samples were re-processed and re-normalized independently to reduce batch effects. The same processing and normalization steps as used in the TD10k cohort were used here. The final processed dataset was used for subsequent association analyses with health outcomes.

### Real DNAm datasets with matched FACS cell counts

*UCLA:* UCLA Immune Assessment Core performed the analysis of immunosenescent cells for 144 whole blood samples, as described previously [96]. Briefly, Total CD3+ T cells, CD4+ T cells, CD8+ T cells, CD19+ B cells, and CD56+/CD16+ NK cells were enumerated in EDTA whole blood with the BD Multitest 6-color TBNK reagent and BD Trucount tubes following the manufacturer’s instructions, acquired on a BD FACSCanto II and analyzed with the BD FACSCanto Software. CD8+ T cell sub setting was performed by staining 50 μl of EDTA whole blood with CD3 FITC, CD8 PerCP, CD28 PE, and CD95 APC (BD) for 10 minutes, followed by BD FACS Lysing used according to the manufacturer’s instructions. At least 10,000 lymphocyte events per sample were acquired and analyzed using DIVA 8.0 software on BD FACSCanto II.

*Koestler:* We used the Illumina 450k dataset from Koestler et al [26] consisting of 6 whole blood (WB) with matched flow-cytometric cell counts. This dataset is available from GEO under accession number GSE77797. DNAm data was normalized and processed as previously described [21].

### Estimation of cell-type fractions

In all cases, given the 12 cell-type DNAm reference matrix for either the Illumina 850k or 450k dataset, we estimated corresponding cell-type fractions using the EpiDISH Bioconductor R-package [21, 97]. Specifically, we ran the *epidish* function with “RPC” as the method and maxit=500.

## Meta-analysis

In each cohort, associations between CTFs and phenotypes was assessed using multivariate linear regression. Covariates generally included age, sex, smoking-status and batch if evidence for batch effects was present and if batch information was available. In general, however, we note that cell-type fractions are relatively robust to batch effects. For specific cohorts were additional covariates were available, multivariate regression models with these additional covariates were also performed. Smoking status was generally treated as ordinal with 0=never-smoker, 1=ex-smoker and 2=current smoker. For each regression we extracted the corresponding Student’s t-test and P-value. To arrive at an overall statistic and P-value for the meta-analysis, we used Stouffer’s method: for each study we transformed the P-value into a normal quantile z-value, using the sign of the statistic to assign the sign of the z-value. We then averaged the z-values over studies using the formula 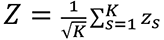 where *K* is the number of studies. From Stouffer’s *Z* value an overall P-value can then be derived using a standard Normal distribution.

### scRNA-Seq dataset of peripheral blood mononuclear cells (PBMCs) from 900 donors

scRNA-Seq data from Yazar et al [42] was downloaded from https://cellxgene.cziscience.com/collections/dde06e0f-ab3b-46be-96a2-a8082383c4a1. Specifically, we downloaded a Seurat object including the cell-by-gene expression matrix as well as associated metadata containing cell type annotation, matched sample IDs, age, sex and UMAP coordinates for 1,248,980 PBMCs from a total of 981 donors. Of the 31 cell types, some were merged as required for comparison with DNAm: CD4+ TCM and CD4+ TEM were regarded as CD4+ T memory cells; CD8+ TCM and CD8+ TEM as CD8+ T memory cells; cells labeled as "NK_CD56bright", "NK Proliferating" and “NK” were regarded as NK cells; CD14+ monocytes and CD16+ monocytes were treated as monocytes; cell types not included in our DNAm reference were regarded as “other”. After this merging, we were left with 9 cell types (not including “other”): approximately 320k CD4+ T memory cells, 259k naïve CD4+ T cells, 177k CD8+ T memory cells, 52k naïve CD8+ T cells, 171k NK, 65k naïve B cells, 30k memory B cells, 51k monocytes and 26k T-regulatory cells. Cell type proportions and differential abundance of these in relation to age and sex were calculated with the propeller method [43] from speckle R-package

### scRNA-Seq dataset of mild and severe Covid-19 cases

scRNA-Seq expression data of bronchoalveolar lavage fluid (BALF) from Liao et al [53] encompassing 3 COVID-19 mild samples, 6 COVID-19 severe samples and 4 healthy controls was obtained from https://cells.ucsc.edu/covid19-balf/exprMatrix.tsv.gz. We also downloaded the corresponding metadata containing annotation for cell types and samples from https://cells.ucsc.edu/covid19-balf/meta.tsv. QC for scRNA-Seq data has been done by Liao et al. by only keeping cells with gene number between 200 and 6000, UMI count > 1000 and mitochondrial gene percentage < 0.1. Cells were annotated after batch effect removal with FindIntegrationAnchors and IntegrateData functions and clustering with FindNeighbors and FindClusters functions from Seurat package [98]. There were clusters mapping to 49417 macrophages, 7716 T cells, 220 B cells, 1607 neutrophils, 3531 epithelial cells, 70 mast cells, 978 mDCs, 1081 NKs, 152 pDCs, 1041 plasma cells, each cluster annotated with signature genes. We used the scRNA-Seq data defined over 23,916 genes and T-cells, B-cells, neutrophils and NKs from 11 samples (3 healthy, 3 mild, 5 severe) for the following analysis, normalizing the data with NormalizeData(normalization.method = "LogNormalize", scale.factor = 10^4^) from Seurat package. Note that among the original 13 samples, one healthy sample (labeled as “HC2” in metadata provided by Liao et al.) and one severe sample (labeled as “S3” in metadata provided by Liao et al.) were removed due to small numbers of T cells profiles in these two samples (<70 cells). T cells were classified as “naïve T cells” if the expression values of LEF1 are non-zero (948 naïve cells), and otherwise classified as “memory/effector T cells” (6653 memory/effector cells). We used the function *propeller(robust = FALSE, trend = FALSE, transform = "asin")* from speckle R-package to calculate for each cell-type their proportions, to perform a variance stabilizing transformation on the proportions and determine whether the differential abundance is statistically significant between non-severe samples and severe samples.

### Health outcome Cox-regression analysis in the MGBB subset of the TD10k cohort

We queried the demographic information (i.e., date of birth, sex and race), health history (i.e., smoking status, alcohol consumption and BMI) and clinical records (i.e., patient diagnosis) of 4386 human subjects from Mass General Brigham (MGB) Biobank [99] and The Research Patient Data Repository (RPDR) databases [100]. Age at the time of sample collection was then calculated accordingly. The vital status (i.e., living/deceased) and date of death were also obtained from MGB Biobank. However, 147 subjects who were recorded as deceased had missing date of death, and they were excluded from the survival analysis of all-cause mortality. We identified other diseases, including type-2 diabetes, chronic obstructive pulmonary disease (COPD), cardiovascular disease (CVD), cancer, and depression, by using relevant ICD-9/10 diagnosis codes (referred to supplement codebook). We defined an incident case as the first diagnosis of a specific health outcome that occurred on the patient’s medical record after the sample collection date. Subjects with a diagnosis code of diseases prior to sample collection were excluded from the survival analysis of that particular disease. We imputed missing data for smoking status, alcohol consumption, and BMI by utilizing the longitudinal records of these variables. Specifically, we used the record closest to the sample collection date for smoking status and alcohol consumption. For BMI, we imputed the missingness as the median value of all BMI records within 6 months around the collection date to balance off the measurement error and temporal variation. Despite imputation, 605 subjects still had missing BMI data and were excluded from the survival analysis when further adjusting for additional risk factors, including BMI. We estimated the hazard ratio of each immune cell type against the health outcomes using Cox proportional hazard regression models with *coxph* function in *survival* R package. The models were adjusted for age, sex and race, and separately again including in addition also smoking status, alcohol consumption and BMI.

### Lasso penalized Cox-regression model predictor of all-cause mortality

Using the same MGBB cohort, the aim was to build a predictor of all-cause mortality using all 12 immune-cell fractions, in addition to age, race, sex, BMI, smoking status and alcohol consumption. We used a penalized (lasso penalty) Cox proportional hazard regression model as implemented in the *glmnet* R-package. Briefly, we divided the dataset up into a 70% training (3591 samples + 302 events) and 30% test (1110 samples + 122 events) set. On the 70% training set we applied an internal 10-fold cross-validation procedure [101], to obtain a risk score for each left-out bag in turn and for each choice of penalty parameter value. The risk scores were then combined across all left-out bags, and the association with all-cause mortality assessed using the C-index. This yielded a curve of how the C-index varies as a function of penalty parameter. We considered the top two models. These models were then tested on the 30% test-test. In all cases, we recorded the Hazard Ratio, C-index and their 95% confidence intervals. P-values of association between the risk scores and all-cause mortality were derived from the one-tailed Chi-square test (1 degree of freedom) as applied to the Cox-score statistic.

## Data Availability

The following DNAm datasets are publicly available from GEO (www.ncbi.nlm.nih.gov/geo/) under accession numbers GSE40279 (Hannum), GSE42861 (LiuRA), GSE50660 (Tsaprouni), GSE106648 (LiuMS), GSE169156 (Song), GSE210255 (HPT-EPIC), GSE210254 (HPT-450k), GSE179325 (Barturen), GSE147740 (Airway), GSE117860 (VACS), GSE87648 (Ventham), GSE84727 (Hannon2), GSE80417 (Hannon1), GSE72680 (Zannas), GSE61151 (Flanagan/FBS), GSE87571 (Johansson), GSE55763 (Lehne). The MRC1946 DNAm data is only available by submitting data requests to; see full policy at http://www.nshd.mrc.ac.uk/data.aspx. Managed access is in place for this study to ensure that use of the data are within the bounds of consent given previously by participants, and to safeguard any potential threat to anonymity since the participants are all born in the same week.

The FCE DNAm data is available from the European Genome Archive (EGA) under accession number EGAS00001005626. The Illumina EPIC DNAm data for the TZH cohort can be viewed at NODE under accession number OEP000260, or directly at https://www.biosino.org/node/project/detail/OEP000260, and accessed by submitting a request for data-access. Data usage shall be in full compliance with the Regulations on Management of Human Genetic Resources in China. All other relevant data supporting the key findings of this study are available within the article and its Supplementary Information files or from the corresponding author upon reasonable request. The TD10k and UCLA datasets are available upon request to TruDiagnostic Inc., in order to protect data privacy of the individuals represented in this cohort.

## Code Availability

The two 12 blood cell-type DNAm reference matrices are available in SI tables S1 and S2, and have also been incorporated into the EpiDISH BioC R-package which includes tools for cell-type fraction estimation: http://www.bioconductor.org/packages/release/bioc/html/EpiDISH.html

## Declarations

### Ethics approval and consent to participate

Only applicable to the TD10k cohort, for which all participants have provided consent to use the collected data for this project.

### Consent for publication

Only applicable to TD10k cohort, for which we have obtained the consent for publication from all participants.

### Availability of data and materials

The following DNAm datasets are publicly available from GEO (www.ncbi.nlm.nih.gov/geo/) under accession numbers GSE40279 (Hannum), GSE42861 (LiuRA), GSE50660 (Tsaprouni), GSE106648 (LiuMS), GSE169156 (Song), GSE210255 (HPT-EPIC), GSE210254 (HPT-450k), GSE179325 (Barturen), GSE147740 (Airway), GSE117860 (VACS), GSE87648 (Ventham), GSE84727 (Hannon2), GSE80417 (Hannon1), GSE72680 (Zannas), GSE61151 (Flanagan/FBS), GSE87571 (Johansson), (GSE55763) Lehne. The MRC1946 DNAm data is only available by submitting data requests to; see full policy at http://www.nshd.mrc.ac.uk/data.aspx. Managed access is in place for this study to ensure that use of the data are within the bounds of consent given previously by participants, and to safeguard any potential threat to anonymity since the participants are all born in the same week. The FCE DNAm data is available from the European Genome Archive (EGA) under accession number EGAS00001005626. The Illumina EPIC DNAm data for the TZH cohort can be viewed at NODE under accession number OEP000260, or directly at https://www.biosino.org/node/project/detail/OEP000260, and accessed by submitting a request for data-access. Data usage shall be in full compliance with the Regulations on Management of Human Genetic Resources in China. All other relevant data supporting the key findings of this study are available within the article and its Supplementary Information files or from the corresponding author upon reasonable request. The TD10k and UCLA datasets are available upon request to TruDiagnostic Inc., in order to protect data privacy of the individuals represented in this cohort. The two 12 blood cell-type DNAm reference matrices are available in SI tables S1 and S2, and have also been incorporated into the EpiDISH BioC R-package which includes tools for cell-type fraction estimation: http://www.bioconductor.org/packages/release/bioc/html/EpiDISH.html

### Authors contributions

QL, VBD, QC, TH, KS performed statistical analyses and contributed to the writing of the manuscript. JMR contributed FACS data. YC, KM, SB and JLS contributed health outcome data. TLM, SV, NE, TZ and RS contributed other data. SCZ helped with software updates. AET, RS and JLS conceived the study. AET wrote the manuscript.

## Acknowledgements

We would like to thank everyone who supports open-access data.

**Description of Additional Files:**

**Additional File 1:** An excel spreadsheet containing all Supplementary Tables.

**Additional File 2:** A pdf Supplementary Information file. Contains all Supplementary Figures and their legends.

